# The YtrBCDEF ABC transporter is involved in the control of social activities in *Bacillus subtilis*

**DOI:** 10.1101/2020.07.24.219923

**Authors:** Martin Benda, Lisa Schulz, Jeanine Rismondo, Jörg Stülke

## Abstract

*Bacillus subtilis* develops genetic competence for the uptake of foreign DNA when cells enter the stationary phase and a high cell density is reached. These signals are integrated by the competence transcription factor ComK which is subject to transcriptional, post-transcriptional and post-translational regulation. Many proteins are involved in the development of competence, both to control ComK activity and to mediate DNA uptake. However, the precise function they play in competence development is often unknown. In this study, we have tested whether proteins required for genetic transformation play a role in the activation of ComK or rather downstream of competence gene expression. While these possibilities could be distinguished for most of the tested factors, two proteins (PNPase and the transcription factor YtrA) are required both for full ComK activity and for the downstream processes of DNA uptake and integration. Further analyses of the role of the transcription factor YtrA for the competence development revealed that the constitutive expression of the YtrBCDEF ABC transporter in the *ytrA* mutant causes the loss of genetic competence. Moreover, constitutive expression of this ABC transporter also interferes with biofilm formation. Since the *ytrGABCDEF* operon is induced by cell wall-targeting antibiotics, we tested the cell wall properties upon overexpression of the ABC transporter and observed an increased thickness of the cell wall. The composition and properties of the cell wall are important for competence development and biofilm formation, suggesting that the increased cell wall thickness as a result of YtrBCDEF overexpression causes the observed phenotypes.

## 1. Introduction

The gram-positive model bacterium *Bacillus subtilis* has evolved many different ways to survive harsh environmental conditions, i. e. it can form highly resistant spores, secrete toxins to kill and cannibalize neighboring cells, form resistant macroscopic biofilms or become competent for transformation (reviewed in (López and Kolter, 2010).

Development of genetic competence is a strategy, which allows bacterial cells to take up foreign DNA from the environment in order to extend the genetic variability of the population. Competence is developed during the transition from exponential to stationary phase of growth as a response to increased cell density and nutrient limitation. In *B. subtilis*, genetic competence is developed in a bistable manner, meaning that only about 10-20% of the cells of a population change their physiological characteristics and become competent for transformation, leaving the rest of the population non-competent in an all or nothing scenario (Haijema et al., 2001; Maamar and Dubnau, 2005). Whether a specific cell becomes competent or not depends on the level of the master regulator ComK (van Sinderen et al., 1995), whose cellular amount is tightly controlled by a complex network of regulators acting on the transcriptional, post-transcriptional as well as on post-translational levels (for a detailed overview see (Maier, 2020).

Transcription of the *comK* gene is controlled by three repressor proteins, Rok, CodY, and AbrB (Serror and Sonenshein, 1996; Hoa et al., 2002; Hamoen et al., 2003), moreover, *comK* transcription is activated by the transcriptional regulator DegU (Hamoen et al., 2000). Another important player for *comK* regulation is Spo0A-P, which controls the levels of the AbrB repressor and additionally supports activation of ComK expression by antagonizing Rok (Mirouze et al., 2012; Hahn et al., 1995). The presence of phosphorylated Spo0A directly links competence to other lifestyles, since Spo0A-P is also involved in pathways leading to sporulation or biofilm formation (Aguilar et al., 2010). When ComK expression reaches a certain threshold, it binds its own promoter region to further increase its own expression, thereby creating a positive feedback loop which leads to full activation of competence (Maamar and Dubnau, 2005; Smits et al., 2005).

ComK levels are also controlled post-transcriptionally by the Kre protein, which destabilizes the *comK* mRNA (Gamba et al., 2015). Post-translational regulation is achieved through the adapter protein MecA, which sequesters ComK and directs it towards degradation by the ClpCP protease (Turgay et al., 1998). During competence, this degradation is prevented by a small protein, ComS, that is expressed in response to quorum sensing (Nakano et al., 1991).

ComK activates expression of more than 100 genes (Berka et al., 2002; Hamoen et al., 2002; Ogura et al., 2002; Boonstra et al., 2020). Whereas a clear role in competence development has been assigned to many of the ComK regulon members, the roles of some ComK-dependent genes remain unclear. Similarly, many single deletion mutant strains were identified as competence deficient, and for many of them the reasons for this deficiency are obvious. However, there are still many single deletion mutants deficient in genetic competence, in which the reason for the loss of competence remains unknown. Typical examples for this are various RNases, namely RNase Y, RNase J1, PNPase or nanoRNase A (Luttinger et al., 1996; Figaro et al., 2013; our unpublished results).Recently, a library of single knock outs of *B. subtilis* genes was screened for various phenotypes, including competence development (Koo et al., 2017). This screen revealed 21 mutants with completely abolished competence. Out of those, 16 are known to be involved in the control of the ComK master regulator, DNA uptake or genetic recombination. However, in case of the other 5 competence-defective strains the logical link to competence is not obvious.

Here, we have focused on some of these factors to investigate their role in genetic competence in more detail. We took advantage of the fact that artificial overexpression of ComK and ComS significantly increases transformation efficiency independently of traditional ComK and ComS regulations (Rahmer et al., 2015). This allows the identification of genes that are involved in competence development due to a function in ComK expression or for other specific reasons downstream of ComK activity. We identified the *ytrGABCDEF* operon as an important player for *B. subtilis* differentiation, since its constitutive expression does not only completely block competence by a so far unknown mechanism, but also affects the proper development of other lifestyles of *B. subtilis.* We discuss the role of thicker cell walls upon overexpression of the proteins encoded by the *ytrGABCDEF* operon as the reason for competence and biofilm defects.

## 2. Materials and Methods

### 2.1. Bacterial strains and growth conditions

The *B. subtilis* strains used in this study are listed in Table 1. Lysogeny broth (LB, Sambrook et al., 1989) was used to grow *E. coli* and *B. subtilis.* When required, media were supplemented with antibiotics at the following concentrations: ampicillin 100 μg ml^−1^ (for *E. coli*) and chloramphenicol 5 μg ml^−1^, kanamycin 10 μg ml^−1^, spectinomycin 250 μg ml^−1^, tetracycline 12.5 μg ml^−1^, and erythromycin 2 μg ml^−1^ plus lincomycin 25 μg ml^−1^ (for *B. subtilis).* For agar plates, 15 g l^−1^ Bacto agar (Difco) was added.

**Table 1.**
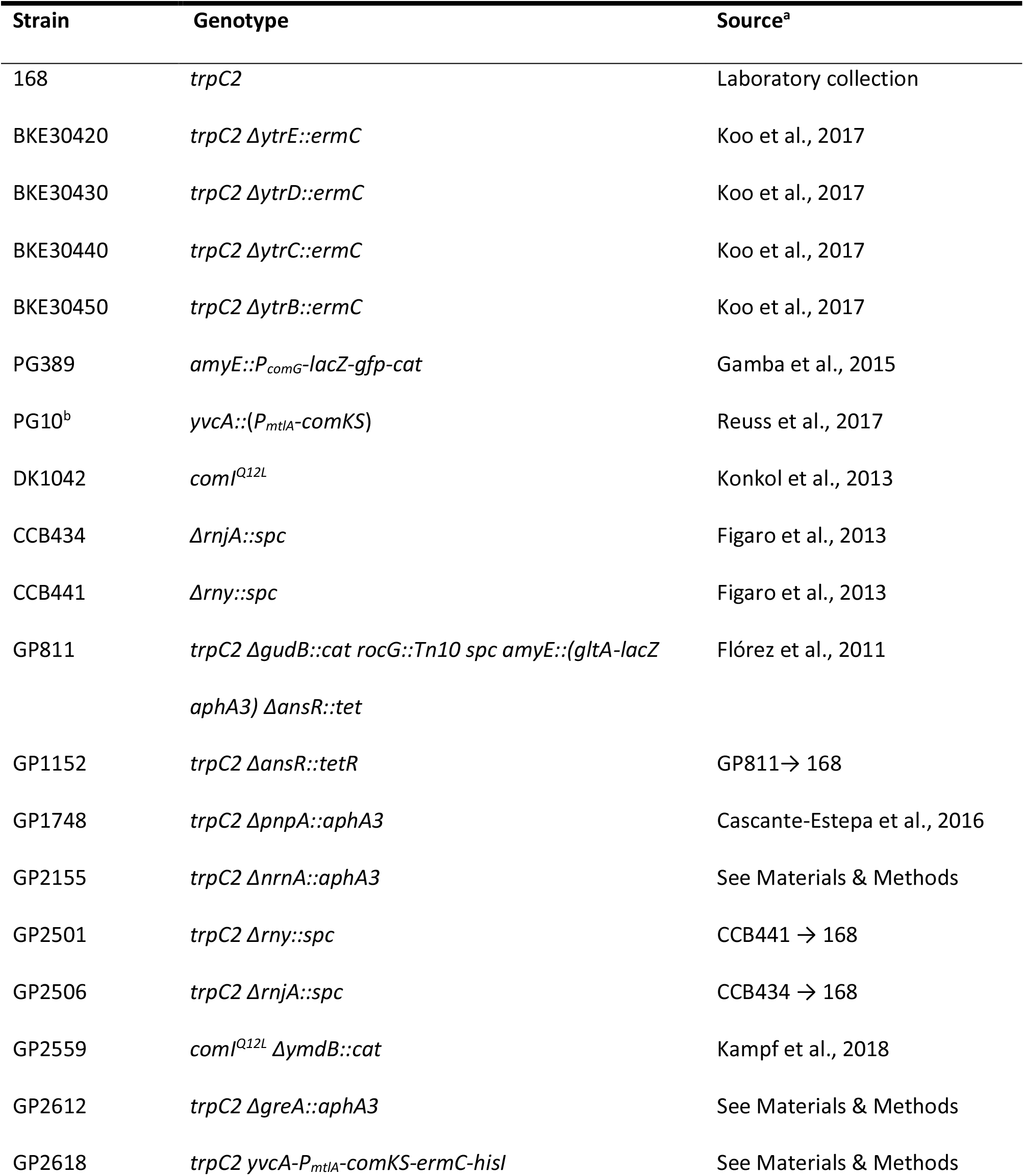
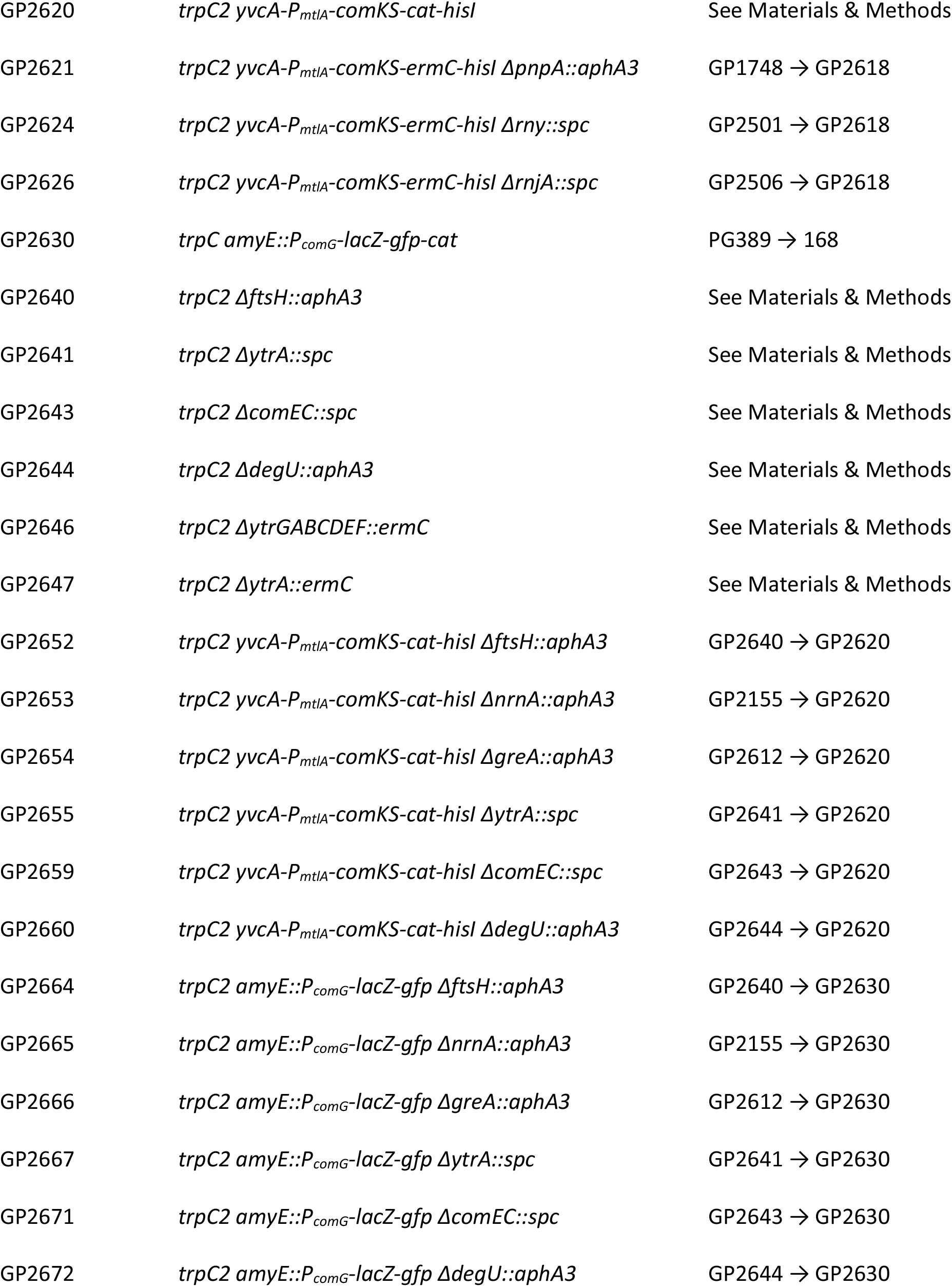
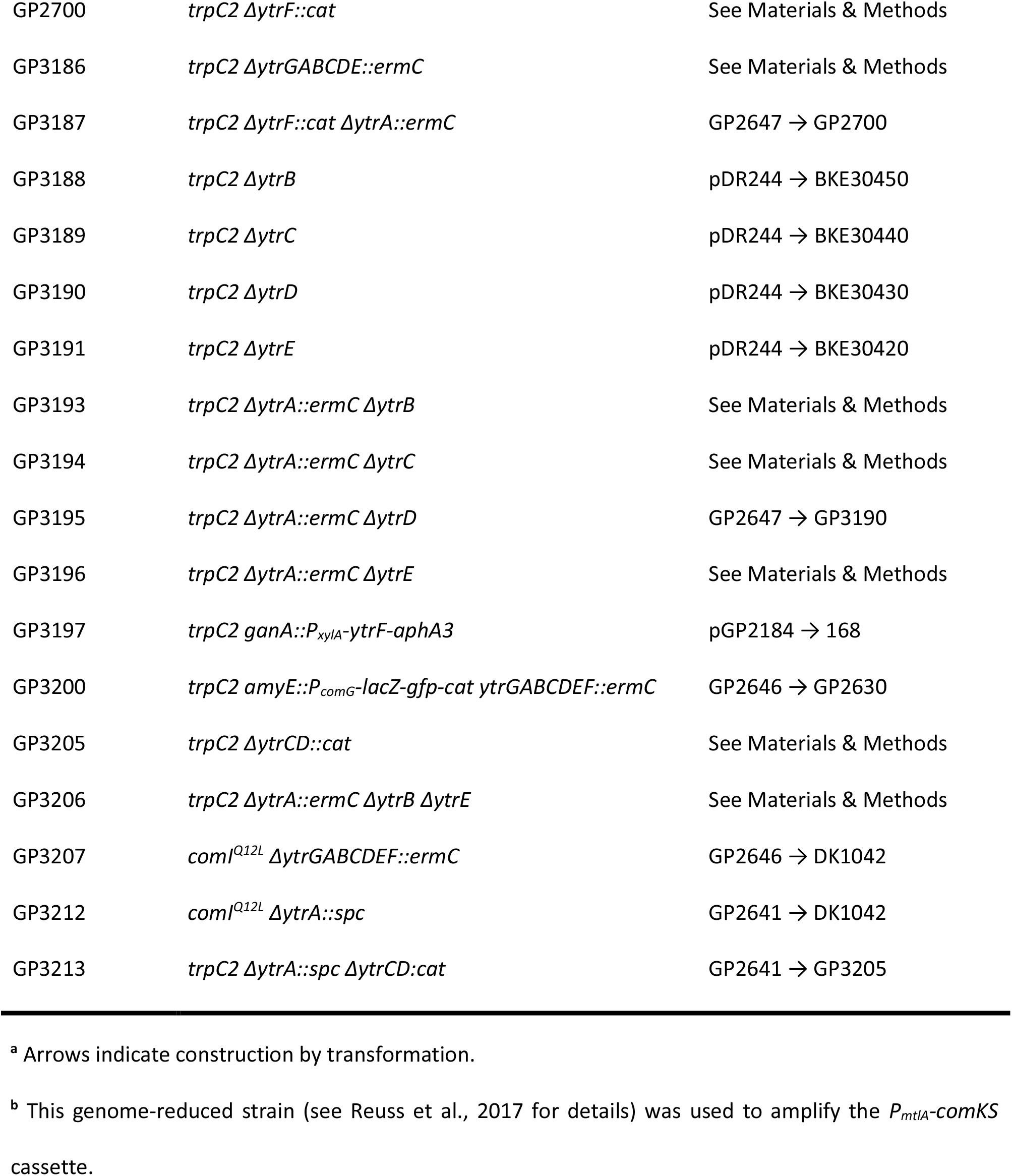
*B. subtilis* strains used in this study.

### 2.2. DNA manipulation and strain construction

S7 Fusion DNA polymerase (Mobidiag, Espoo, Finland) was used as recommended by the manufacturer. DNA fragments were purified using the QIAquick PCR Purification Kit (Qiagen, Hilden, Germany). DNA sequences were determined by the dideoxy chain termination method (Sambrook et al., 1989). Chromosomal DNA from *B. subtilis* was isolated using the peqGOLD Bacterial DNA Kit (Peqlab, Erlangen, Germany) and plasmids were purified from *E. coli* using NucleoSpin Plasmid Kit (Macherey-Nagel, Düren, Germany). Deletion of the *degU, comEC, ftsH, greA, ytrA, nrnA*, and *ytrF* genes as well as *ytrCD, ytrG-ytrE*, and *ytrGABCDEF* regions was achieved by transformation with PCR products constructed using oligonucleotides (see Table S1) to amplify DNA fragments flanking the target genes and intervening antibiotic resistance cassettes as described previously (Guérout-Fleury et al., 1995; Wach, 1996, Youngman, 1990). The identity of the modified genomic regions was verified by DNA sequencing. To construct the strains (GP2618 and GP2620) harbouring the *PmtlA-comKS* cassette coupled to the antibiotic resistance gene, we have first amplified the *PmtlA-comKS* from the strain PG10 (Reuß et al., 2017) as well as the resistance genes from pDG646 and pGEM-cat, respectively (Youngman, 1990; Guérout-Fleury et al., 1995) and the genes flanking the intended integration site, i. e. *yvcA* and *hisI* from *B. subtilis* 168. Subsequently, those DNA fragments were fused in another PCR reaction thanks to the overlapping primers. The final product was used to transform *B. subtilis* 168. Correct insertion was verified by PCR amplification and sequencing. Markerless deletions of *ytrB, ytrC, ytrD* and *ytrE* genes were performed using pDR244 plasmid as described (Koo et al., 2017). In short, strains BKE30450, BKE30440, BKE30430 and BKE30420 were transformed with plasmid pDR244 and transformants were selected on LB agar plates supplemented with spectinomycin at 30°C. Transformants were then streaked on plain LB agar plates and incubated at 42°C to cure the plasmid, which contains a thermo-sensitive origin of replication. Single colonies were then screened for spectinomycin and erythromycin/lincomycin sensitivity. Markerless deletion was confirmed by PCR with primers flanking the deletion site. Created strains GP3188, GP3189, GP3190 and GP 3191 were used for subsequent deletion of the *ytrA* gene. This was done either by transformation with PCR product as described above or by transformation with genomic DNA of the *ytrA* deletion strain (in case of GP3195 construction). Deletion of the *ytrA* gene and preservation of selected markerless deletions were confirmed via PCR. To construct GP3206, PCR product containing erythromycin resistance in place of *ytrA* and *ytrB* genes was amplified from GP3193 and transformed to GP3191.

### 2.3. Transformation of *B. subtilis* strains

Transformation experiments were conducted based on the two-step protocol as described previously (Kunst and Rapoport, 1995). Briefly, cells were grown at 37°C at 200 rpm in 10 ml MNGE medium containing 2% glucose, 0.2% potassium glutamate, 100 mM potassium phosphate buffer (pH 7), 3.4 mM trisodiumcitrate, 3 mM MgSO_4_, 42 μM ferric ammonium citrate, 0.24 mM L-tryptophan and 0.1% casein hydrolysate. During the transition from exponential to stationary phase, the culture was diluted with another 10 ml of MNGE medium (without casein hydrolysate) and incubated for 1 h at 37°C with shaking. In case of strain GP3187, 0.5% xylose was added to both media. Afterwards, 250 ng of chromosomal DNA was added to 400 μl of cells and incubated for 30 minutes at 37°C. One hundred microliter of Expression mix (2.5% yeast extract, 2.5% casein hydrolysate, 1.22mM tryptophan) was added and cells were allowed to grow for 1h at 37°C, before spreading onto selective LB plates containing appropriate antibiotics.

Transformation of strains harboring *comK* and *comS* expressed from the mannitol inducible promotor (P_*mtlA*_) was performed based on (Rahmer et al., 2015). Briefly, an overnight culture was diluted in 5 ml LB to an initial OD_600_ of 0.1 and incubated at 37°C at 200 rpm. After 90 minutes incubation, 5 ml of fresh LB containing mannitol (1%) and MgCl_2_ (5 mM) were added and the bacterial culture was incubated for an additional 90 minutes. The cells were then pelleted by centrifugation for 10 minutes at 2,000 g and the pellet was re-suspended in the same amount of fresh LB medium, 1 ml aliquots were distributed into 1.5 ml reaction tubes and 250 ng of chromosomal DNA was added to each of them. The cell suspension was incubated for 1 h at 37°C and transformants were selected on LB plates as described above.

### 2.4. Plasmid construction

All plasmids used in this study are listed in Table 2. *Escherichia coli* DH5α (Sambrook et al., 1989) was used for plasmid constructions and transformation using standard techniques (Sambrook et al., 1989). To express the *B. subtilis* protein YtrF under the control of a xylose inducible promotor, we cloned the *ytrF* gene into the backbone of pGP888 via the XbaI and KpnI sites (Diethmaier et al., 2011).

**Table 2.** Plasmids used in this study.

| Plasmid | Relevant Characteristics | Primers | Reference |
| --- | --- | --- | --- |
| pDR244 | *cre* + Ts origin | - | Koo et al., 2017 |
| pGEM-cat | Amplification of the cat cassette | - | Youngman, 1990 |
| pDG646 | Amplification of the ermC cassette | - | Guérout-Fleury et al., 1995 |
| pDG780 | Amplification of the aphA3 cassette | - | Guérout-Fleury et al., 1995 |
| pDG1726 | Amplification of the spc cassette | - | Guérout-Fleury et al., 1995 |
| pGP888 | *ganA*::P<sub>xyIA</sub>; aphA3 | - | Diethmaier et al., 2011 |
| pGP2184 | pGP888-*ytrF* | MB186/MB187 | This study |

### 2.5. Biofilm assay

To analyse biofilm formation, selected strains were grown in LB medium to an OD_600_ of about 0.5 to 0.8 and 10 μl of the culture were spotted onto MSgg agar plates (Branda et al., 2001). Plates were incubated for 3 days at 30°C.

### 2.6. Fluorescence microscopy

For fluorescence microscopy imaging, *B. subtilis* cultures were grown in 10 ml MNGE medium till the transition from exponential to stationary phase and then diluted with another 10 ml of MNGE medium as described for the transformation experiments (see section 2.3). 5 μl of cells were pipetted on microscope slides coated with a thin layer of 1% agarose and covered with a cover glass. Fluorescence images were obtained with the AxioImager M2 fluorescence microscope, equipped with digital camera AxioCam MRm and AxioVision Rel 4.8 software for image processing and an EC Plan-NEOFLUAR 100X/1.3 objective (Carl Zeiss, Göttingen, Germany). Filter set 38 (BP 470/40, FT 495, BP 525/50; Carl Zeiss) was applied for GFP detection. Ratio of GFP expressing cells to the total number of cells was determined by manual examination from three independent randomly selected pictures originated from at least two independent growth replicates.

### 2.7. Transmission electron microscopy

To examine cell wall thickness of *B. subtilis* strains, cells were prepared for Transmission Electron Microscopy (TEM) as previously described (Rincón-Tomas et al., 2020). An overnight culture was inoculated to an OD_600_ of 0.05 in 30 ml MNGE medium and grown to an OD_600_ of 0.6 ± 0.1 at 37°C and 200 rpm. Cells were centrifuged for 10 minutes at 4,000 rpm to obtain a 100 μl cell pellet, which was then washed twice in phosphate-buffered saline (PBS, 127 mM NaCl, 2.7 mM KCl, 10 mM Na2HPO4, 1.8 mM KH_2_PO_4_, pH 7.4) and fixed overnight in 2.5% (w/v) glutaraldehyde at 4°C. Cells were then mixed with 1.5% (w/v, final concentration) molten Bacto-Agar (in PBS) and the resulting agar block was cut to pieces of 1 mm^3^. A dehydration series was performed (15% aqueous ethanol solution for 15 minutes, 30%, 50%, 70% and 95% for 30 minutes and 100% for 2 x 30 minutes) at 0°C, followed by an incubation step in 66% LR white resin mixture (v/v, in ethanol) (Plano, Wletzlar, Germany) for 2 hours at room temperature and embedment in 100% LR-White solution overnight at 4°C. One agar piece was transferred to a gelatin capsule filled with fresh LR-white resin, which was subsequently polymerized at 55°C for 24 hours. A milling tool (TM 60, Fa. Reichert & Jung, Vienna, Austria) was used to shape the gelatin capsule into a truncated pyramid. An ultramicrotome (Reichert Utralcut E, Leica Microsystems, Wetzlar, Germany) and a diamond knife were used to obtain ultrathin sections (80 nm) of the samples. The resulting sections were mounted onto mesh specimen Grids (Plano, Wetzlar, Germany) and stained with 4% (w/v) uranyl acetate solution (pH 7.0) for 10 minutes. Microscopy was performed in a Joel JEM 1011 transmission electron microscope (Joel Germany GmbH, Freising, Germany) at 80 kV. Images were taken at a magnification of 30,000 and recorded with a Gatan Orius SC1000 CCD camera (Gatan, Munich, Germany). For each replicate, 20 cells were photographed and cell wall thickness was measured at three different locations using ImageJ software (Rueden et al., 2017).

## 3. Results

### 3.1. ComK-dependent and –independent functions of proteins required for the development of genetic competence

Genetic work with *B. subtilis* is facilitated by the development of genetic competence, a process that depends on a large number of factors. While the specific contribution of many proteins to the development of competence is well understood, this requirement has not been studied for many other factors. In particular, several RNases (RNase Y, RNase J1, PNPase and nanoRNase A) are required for competence, and the corresponding mutants have lost the ability to be become naturally competent (Luttinger et al., 1996; Figaro et al., 2013; our unpublished results). We are interested in the reasons for the loss of competence in these mutant strains, as well as in other single gene deletion mutants which are impaired in the development of natural competence for unknown reasons (Koo et al., 2017). Therefore, we first tested the roles of the aforementioned RNases (encoded by the *rny*, *rnjA, pnpA, and nrnA* genes) as well as of the transcription elongation factor GreA, the metalloprotease FtsH and the transcription factor YtrA (Koo et al., 2017) for the development of genetic competence. For this purpose, we compared the transformation efficiencies of the corresponding mutant strains to that of a wild type strain. We have included two controls to all experiments, i. e. *comEC* and *degU* mutants. Both mutants have completely lost genetic competence, however for different reasons. The ComEC protein is directly responsible for the transport of the DNA molecule across the cytoplasmic membrane. Loss of ComEC blocks competence, but it should not affect the global regulation of competence development and expression of other competence factors (Draskovic and Dubnau, 2005). In contrast, DegU is a transcription factor required for the expression of the key regulator of competence, ComK, and thus indirectly also for the expression of all other competence genes (Hamoen et al., 2000; Shimane and Ogura, 2004). Our analysis confirmed the significant decrease in transformation efficiency for all tested strains (see Table 3). For five out of the seven strains, as well as the two control strains competence was abolished completely, whereas transformation of strains GP2155 (*ΔnrnA*) and GP1748 (*ΔpnpA*) was possible, but severely impaired as compared to the wild type strain. This result confirms the implication of these genes in the development of genetic competence.

**Table 3.** Effect of gene deletions on the development of genetic competence in dependence of the competence transcription factor ComK^a^.

|  | Wild type | *P<sub>mtlA</sub>-comKS* |
| --- | --- | --- |
| Mutant | Colonies per µg of DNA |  |
| Wild type | 138,600 ± 17,006 | 47,952 ± 8,854 |
| *ΔdegU* | 0 ± 0 | 60,853 ± 13,693 |
| *ΔcomEC* | 0 ± 0 | 0 ± 0 |
| *ΔnrnA* | 1,689 ± 316 | 34,933 ± 6,378 |
| *ΔftsH* | 0 ± 0 | 0 ± 0 |
| *ΔgreA* | 0 ± 0 | 0 ± 0 |
| *Δrny* | 0 ± 0 | 0 ± 0 |
| *ΔrnjA* | 0 ± 0 | 0 ± 0 |
| *ΔpnpA* | 17 ± 6 | 293 ± 19 |
| *ΔytrA* | 0 ± 0 | 467 ± 278 |

<sup>a</sup> Cells were transformed with chromosomal DNA of strain GP1152 harboring a tetracycline resistance marker as described in Material and Methods.

The proteins that are required for genetic competence might play a more general role in the control of expression of the competence regulon (as known for the regulators that govern ComK expression and stability, e. g. the control protein DegU), or they may have a more specific role in competence development such as the control protein ComEC. To distinguish between these possibilities, we introduced the mutations into a strain that allows inducible overexpression of the *comK* and *comS* genes. The overexpression of *comK* and *comS* allows transformation in rich medium and hence facilitates the transformation of some competence mutants (Rahmer et al., 2015). For this purpose, we first constructed strains that contain mannitol inducible *comK* and *comS* genes fused to resistance cassettes (GP2618 and GP2620, for details see Materials and Method). Subsequently, we deleted our target genes in this genetic background and assayed transformation efficiency after induction of *comKS* expression (for details see Materials and Method). In contrast to the strain with wild type *comK* expression, the transformation efficiency of the *degU* mutant was now similar to the isogenic wild type strain. This suggests that DegU affects competence only by its role in *comK* expression and that DegU is no longer required in the strain with inducible *comKS* expression. In contrast, the *comEC* mutant was even in this case completely non-competent, reflecting the role of the ComEC protein in DNA uptake (see Table 3). Of the tested strains, only the *nrnA* mutant showed a transformation efficiency similar to that of the isogenic control strain with inducible *comKS* expression. This observation suggests that nanoRNase A might be involved in the control of *comK* expression. In contrast, the *ftsH, greA, rny* and *rnjA* mutants did not show any transformants even upon *comKS* overexpression, indicating that the corresponding proteins act downstream of *comK* expression. Finally, we have observed a small but reproducible restoration of competence in case of the *pnpA* and *ytrA* mutants. This finding is particularly striking in the case of the *ytrA* mutant, since this strain did not yield a single transformant in the 168 background (see Table 3). However, the low number of transformants obtained with *pnpA* and *ytrA* mutants as compared to the isogenic wild type strain suggests that PNPase and the YtrA transcription factor play as well a role downstream of *comK.*

ComK activates transcription of many competence genes including *comG* (van Sinderen et al., 1995). Therefore, as a complementary approach to further verify the results shown above, we decided to assess ComK activity using a fusion of the *comG* promoter to a promoterless GFP reporter gene (Gamba et al., 2015). For this purpose, we deleted the selected genes in the background of strain GP2630 containing the P*_comG_-gfp* construct. We grew the cells in competence inducing medium using the two-step protocol as we did for the initial transformation experiment. At the time point, when DNA would be added to the cells during the transformation procedure, we assessed *comG* promoter activity in the cells using fluorescence microscopy. Since expression of ComK and thus also activation of competence takes place only in sub-population of cells (Smits et al., 2005), we determined the ratio of *gfp* expressing cells as an indication of ComK activity for each of the strains (see Table 4). Since RNase mutants tend to form chains, thus making it difficult to study florescence in individual cells, we did not include the RNase mutants for this analysis.

**Table 4.**
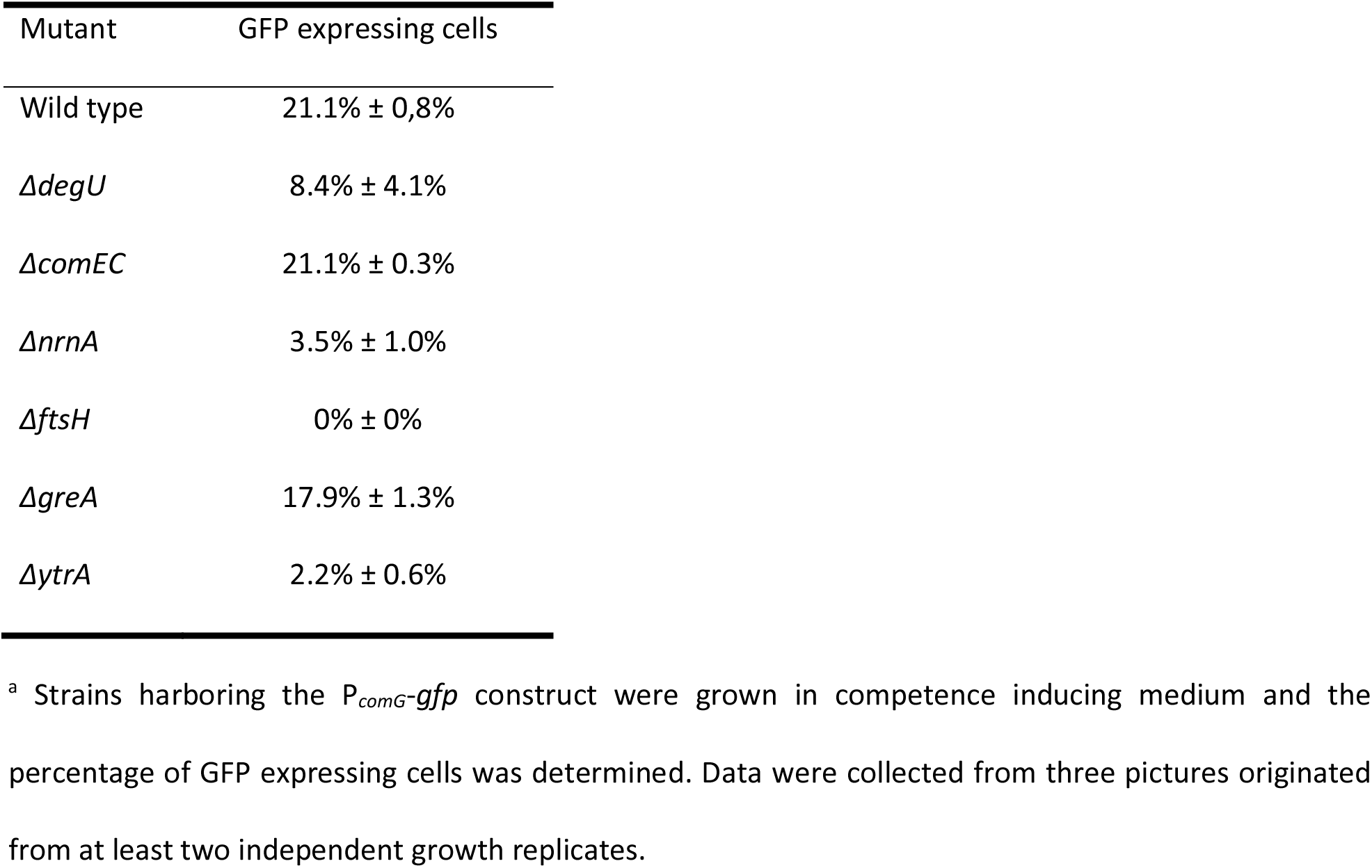
Effect of gene deletions on the activity of the competence transcription factor ComK as studied by the percentage of cells expressing a P*_comG_-gfp* transcriptional fusion^a^.

| Mutant | GFP expressing cells |
| --- | --- |
| Wild type | 21.1% ± 0,8% |
| *ΔdegU* | 8.4% ± 4.1% |
| *ΔcomEC* | 21.1% ± 0.3% |
| *ΔnrnA* | 3.5% ± 1.0% |
| *ΔftsH* | 0% ± 0% |
| *ΔgreA* | 17.9% ± 1.3% |
| *ΔytrA* | 2.2% ± 0.6% |

<sup>a</sup> Strains harboring the $P_{comG}$ -*gfp* construct were grown in competence inducing medium and the percentage of GFP expressing cells was determined. Data were collected from three pictures originated from at least two independent growth replicates.

In the wild type strain GP2630, about 20% of the cells expressed GFP, and similar numbers were obtained for the control strain lacking ComEC, which is not impaired in *comK* and subsequent *comG* expression. In contrast, the control strain lacking DegU showed decreased amount of GFP expressing cells as compared to the wild type, which reflects the role of DegU in the activation of *comK* expression. In agreement with our previous finding that nanoRNase A affects ComK activity, only about 3% of *nrnA* mutant cells showed expression from P*_comG_-gfp* For the *ftsH* mutant, we did not find any single cell expressing GFP. This is striking since our previous results suggested that ComK expression is not the cause of competence deficiency in this case. For the strain lacking GreA, we observed similar rates of GFP expressing cells as in the wild-type strain, indicating that ComK activation is not the problem that causes loss of competence. Finally, we have observed significantly decreased ratio of GFP producing cells in case of the *ytrA* deletion mutant.

Taken together we have discovered that *nrnA* coding for nanoRNase A (Mechold et al., 2007) plays a so far undiscovered role in the regulation of *comK.* In contrast, the GreA transcription elongation factor is required for competence development in steps downstream of *comK* expression. FtsH and YtrA seem to play a dual role in the development of genetic competence. On one hand, they are both required for ComK activity but on the other hand, they have a ComK-independent function. The *ytrA* gene encodes a transcription factor with a poorly studied physiological function (Salzberg et al., 2011). Therefore, we focused our further work on understanding the role of this gene in development of genetic competence.

### 3.2. Overexpression of the YtrBCDEF ABC transporter inhibits genetic competence

The *ytrA* gene encodes a negative transcription regulator of the GntR family, which binds to the inverted repeat sequence AGTGTA-13bp-TACACT (Salzberg et al., 2011). In the *B. subtilis* genome, this sequence is present in front of two operons, its own operon *ytrGABCDEFG* and *ywoBCD.* The deletion of *ytrA* leads to an overexpression of these two operons (Salzberg et al., 2011). It is tempting to speculate that overexpression of one of these operons is the cause for the loss of competence in the *ytrA* mutant. To test this hypothesis, we constructed strain GP2646, which lacks the complete *ytrGABCDEF* operon. Next, we assayed the genetic competence of this strain. This revealed that although deletion of *ytrA* fully blocks genetic competence, the strain lacking the whole operon is transformable in similar rates as the wild type strain 168 (Table 5). We conclude that overexpression of the *ytrGABCDEF* operon causes the loss of competence in the *ytrA* mutant strain. In addition, we tested ComK activity in the mutant lacking the operon, using the expression of the *PcomG-gfp* fusion as a readout. As observed for the wild type, about 20% of the mutant cells expressed *comG*, indicating that ComK is fully active in the mutant, and that the reduced activity in the *ytrA* mutant results from the overexpression of the operon (data not shown). Initially we also attempted deleting the *ywoBCD* operon, however we failed to construct such a strain in several experiments. As we have already discovered that the overexpression of the *ytr* operon causes the loss of competence in the *ytrA* mutant, we decided not to continue with this second YtrA-controlled operon.

**Table 5.** Effect of gene deletions in the *ytrGABCDEF* operon on the development of genetic competence^a^.

| Mutant | Colonies per µg of DNA |
| --- | --- |
| Wild type | 138,600 ± 17,006 |
| *ΔytrGABCDEF* | 114,733 ± 14,408 |
| *ΔytrA* | 0 ± 0 |
| *ΔytrAB* | 0 ± 0 |
| *ΔytrAC* | 0 ± 0 |
| *ΔytrAD* | 24 ± 2 |
| *ΔytrAE* | 137 ± 51 |
| *ΔytrAF* | 10,180 ± 549 |
| *P<sub>xyI</sub>-ytrF* | 137,533 ± 26,595 |
| *ΔytrGABCDE* | 108,467 ± 14,836 |
| *ΔytrABE* | 309 ± 88 |
| *ΔytrACD* | 45,467 ± 10,799 |

<sup>a</sup> Cells were transformed with chromosomal DNA of strain GP1152 harboring a tetracycline resistance marker as described in Material and Methods.

The *ytr* operon consist of seven genes (see Fig 1A). Five proteins encoded by this operon (YtrB, YtrC, YtrD, YtrE and YtrF) are components of a putative ABC transporter (see Fig 1B), which was suggested to play a role in acetoin utilization (Quentin et al., 1999; Yoshida et al., 2000). YtrB and YtrE are supposed to be the nucleotide binding domains, YtrC and YtrD the membrane spanning domains and YtrF the substrate binding protein. Finally, another open reading frame called *ytrG*, encodes a peptide of 45 amino acids which is unlikely to be part of the ABC transporter (Salzberg et al., 2011). The expression of the *ytr* operon is usually kept low due to transcriptional repression exerted by YtrA. This repression is naturally relieved only in response to several lipid II-binding antibiotics or during cold-shock (Salzberg et al., 2011; Wenzel et al., 2012, Beckering et al., 2002).

**FIG 1.**
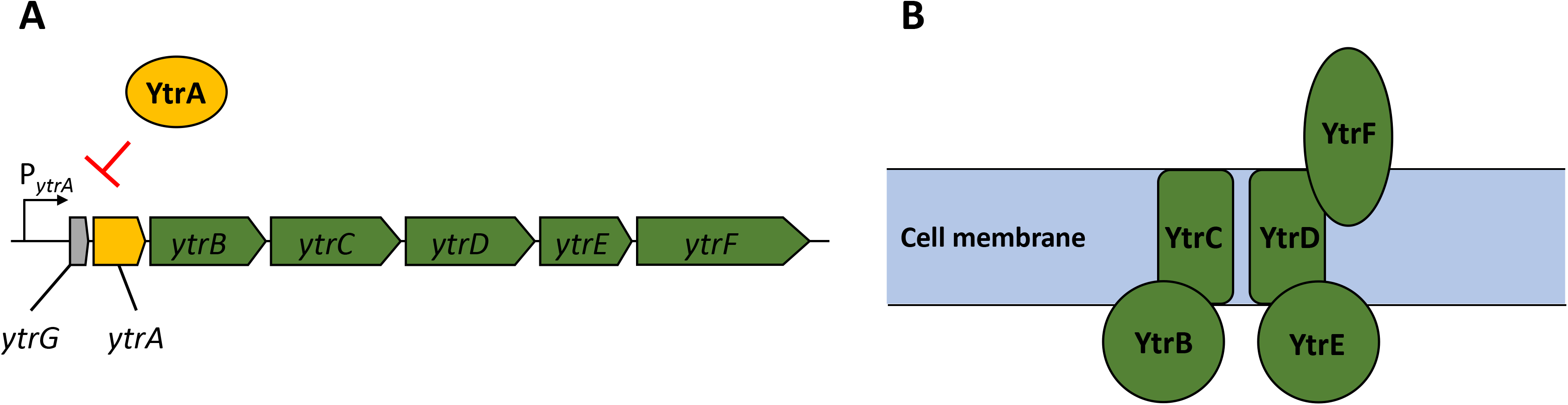
Genetic organization of the *ytrGABCDEF* operon and organization of the putative ABC transporter. **(A)** Reading frames are depicted as arrows with respective gene names. Green arrows indicate proteins suggested to form the ABC transporter; the yellow arrow indicates the gene coding for the repressor YtrA and the grey arrow indicates the small open reading frame called *ytrG.* The map was constructed based on information provided in Salzberg et al. (2011) **(B)** Organization of the putative ABC transporter YtrBCDEF as suggested by Yoshida et al. (2000). YtrB and YtrE are nucleotide binding proteins, YtrC and YtrD membrane spanning proteins and YtrF is a solute binding protein. The role and localization of the YtrG peptide remain elusive.

To test the involvement of the individual components of the putative YtrBCDEF ABC transporter in the development of genetic competence, we constructed double mutants of *ytrA* together with each one of the other genes of the operon, i.e. *ytrB, ytrC, ytrD, ytrE* and *ytrF.* The results (Table 5) revealed that most of the double mutants are deficient in genetic transformation, as observed for the single *ytrA* mutant GP2647. However, strain GP3187 with deletions of *ytrA* and *ytrF* but still overexpressing all the other parts of the transporter, had partially restored competence. We conclude that the YtrF protein is the major player for the loss of competence in the overexpressing strain.

To further test the role of YtrF overexpression for the loss of competence, we used two different approaches. First, we constructed a strain with artificial overexpression of *ytrF* from a xylose inducible promoter (GP3197) and second, we created a strain with deletion of all other components (*ytrGABCDEF*) of the operon, leaving only constitutively expressed *ytrF* (GP3186). In contrast to our expectations, competence was not blocked in any of the two strains, suggesting that increased presence of YtrF protein alone is not enough to block the competence and that YtrF might need assistance from the other proteins of the putative transporter for its full action/proper localization. The *ytr* operon encodes two putative nucleotide binding proteins (YtrB and YtrE) and two putative membrane spanning proteins (YtrC, YtrD), whereas YtrF is the only solute binding protein that interacts with the transmembrane proteins. Therefore, we hypothesized that YtrF overexpression might only block genetic competence if the protein is properly localized in the membrane via YtrC and YtrD. To check this possibility, we constructed strains GP3206 and GP3213 lacking YtrA and the nucleotide binding proteins or the membrane proteins, respectively, and tested their transformability. Strain GP3206 showed very few transformants, suggesting that the presence of nucleotide binding proteins is not important to block competence. In contrast, strain GP3213 gave rise to many transformants. We thus conclude that the overexpression of the solute binding protein YtrF in conjunction with the membrane proteins YtrC and YtrD is responsible for the block of competence indicating that indeed the proper function of YtrF, which depends on YtrC and YtrD, is crucial for the phenotype.

### 3.3. Overexpression of the *ytrGABCDEF* operon leads to defect in biofilm formation

*B. subtilis* can employ various lifestyles which are tightly interconnected through regulatory proteins (Lopez et al., 2009). Therefore, we anticipated that the overexpression of YtrF might also affect other lifestyles of *B. subtilis.* Indeed, it was previously shown that the *ytrA* mutant has a reduced sporulation efficiency (Koo et al., 2017). We thus decided to examine the effect of the *ytrA* deletion on biofilm formation. To that end, we first deleted the *ytrA* gene or the whole *ytrGABCDEF* operon from the biofilm-forming strain DK1042 (Konkol et al., 2013). We then tested the biofilm formation of the resulting strains on biofilm inducing MSgg agar (Branda et al., 2001). As expected, the wild type strain DK1042 formed structured colonies that are indicative of biofilm formation. In contrast, the negative control GP2559 (a *ymdB* mutant that is known to be defective in biofilm formation, Kampf et al., 2018) formed completely smooth colonies. The biofilm formed by the *ytrA* mutant GP3212 was less structured, more translucent and with only some tiny wrinkles on its surface, indicating that biofilm formation was inhibited but not fully abolished upon loss of YtrA. In contrast, strain GP3207 lacking the complete *ytrGABCDEF* operon formed biofilm indistinguishable from the one of the parental strain DK1042 (see Fig. 2). This observation suggests that overexpression of components of the Ytr ABC transporter interferes with biofilm formation.

**FIG 2.**
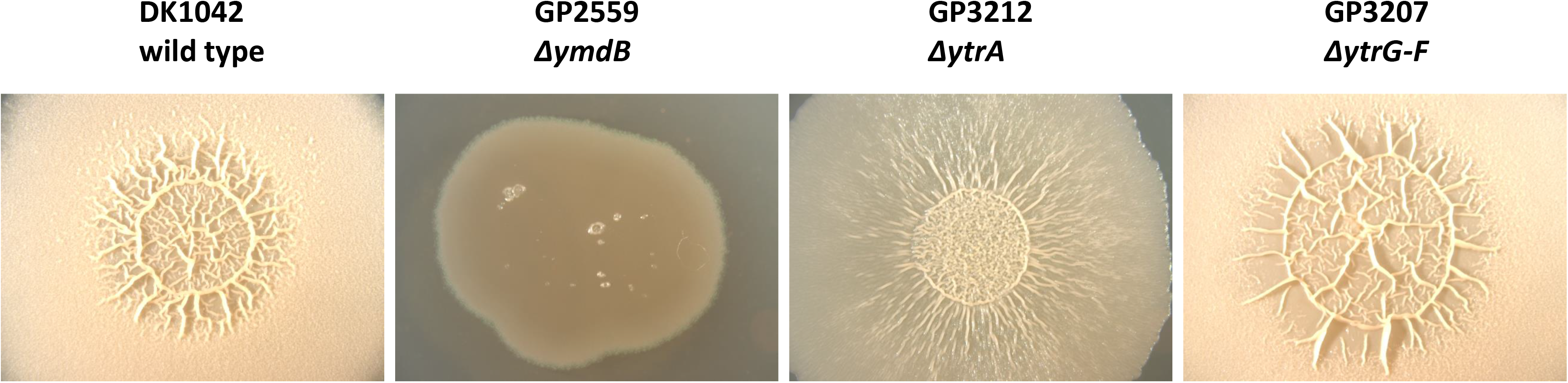
Biofilm formation is affected by the *ytrA* deletion. Biofilm formation was examined in the wild type strain DK1042 and respective deletion mutants of *ymdB* (G2559), *ytrA* (GP3212) and *ytrGABCDEF* (GP3207). The biofilm assay was performed on MSgg agar plates as described in Material and Methods. The plates were incubated for 3 days at 30°C. All images were taken at the same magnification.

### 3.4. Overexpression of the *ytr* operon increases cell wall thickness

In previous experiments, we have shown that the expression of the *ytr* operon interferes with the development of genetic competence and biofilm formation due to the activity of the solute binding protein YtrF. However, it remains unclear why competence and biofilm formation are abolished. The *ytr* operon is repressed under standard conditions by the YtrA transcription regulator and this repression is naturally relieved only upon exposure to very specific stress conditions, mainly in response to cell wall targeting antibiotics and cold shock (Beckering et al., 2002; Cao et al., 2002; Mascher et al., 2003; Salzberg et al., 2011; Nicolas et al., 2012; Wenzel et al., 2012). The possible link between antibiotic resistance, genetic competence, and biofilm formation is not apparent, however, cell wall properties might provide an answer. Indeed, it has been shown that wall teichoic acids, the uppermost layer of the cell wall, are important for DNA binding during the process of transformation and biofilm formation (Mirouze et al., 2018; Bucher et al., 2015; Zhu et al., 2018).

To test the hypothesis that overexpression of the putative ABC transporter encoded by the *ytrGABCDEF* operon affects cell wall properties of the *B. subtilis* cells, we decided to compare the cell morphology of the wild type and the *ytrA* mutant as well as the *ytrGABCDEF* mutant lacking the complete operon by transmission electron microscopy. While the wild type strain showed an average cell wall thickness of 21 nm, which is agreement with previous studies (Beveridge and Murray, 1979), the *ytrA* (GP2647) mutant showed a significant increase in cell wall thickness with an average of 31 nm. In contrast, such an increase was not observed for the whole operon mutant (GP2646) that had an average cell wall thickness of 23 nm (see Fig 3). These observations are in excellent agreement with the hypothesis that the overexpression of the YtrBCDEF ABC transporter affects cell wall properties and thereby genetic competence and biofilm formation.

**FIG 3.**
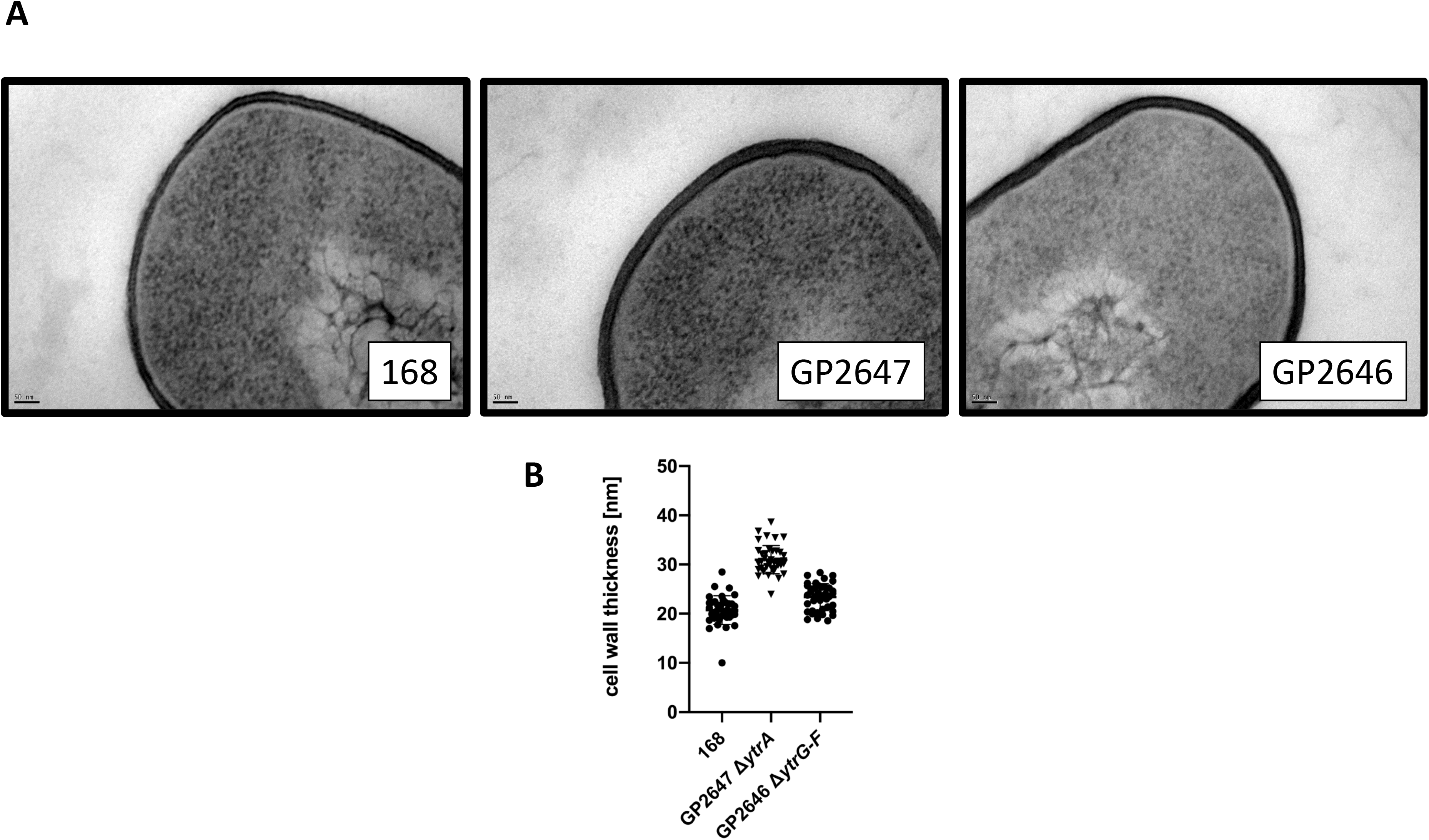
The *ytrA* mutant has thicker cell walls. **(A)** Shown are representative transmission electron microscopy images of the wild type strain 168, the *ytrA* mutant (GP2647) and the whole operon *ytrGABCDEF* mutant (GP2646). **(B)** The graph shows the cell wall thickness of 40 individual measurements from two growth replicates as described in Material and Methods.

## 4. Discussion

In this work we have shown that overexpression of the *ytrGABCDEF* operon, coding for a so far uncharacterized ABC transporter, completely blocks the development of genetic competence and interferes with biofilm formation in *B. subtilis.* This block is mediated by the solute binding protein YtrF in cooperation with at least one membrane spanning protein (YtrC or YtrD) that are required for correct function of YtrF. The overexpression of the YtrBCDEF ABC transporter is the reason for the loss of competence of an *ytrA* regulator mutant that had been observed in a previous genome-wide study (Koo et al., 2017). Based on its expression pattern, the *ytr* operon was described as a reporter for glycopeptide antibiotics, such as vancomycin or ristocetin (Hutter et al., 2004) and later also for other antibiotics that interfere with the lipid II cycle, such as nisin (Wenzel et al., 2012). Whether this induction of *ytrGABCDEF* expression leads to an increased resistance towards those antibiotics is not clear, but recent results indicate that it does at least in case of nisin (J. Bandow, personal communication).

Based on the partial restoration of genetic competence of the *ytrA* mutant upon ComKS overexpression, one might expect that the loss of YtrA and the concomitant overexpression of the ABC transporter somehow interferes with competence development upstream of ComK activation. However, competence is developed in an all or nothing scenario, and cells in which the ComK levels reach a certain threshold should become competent (Haijema et al., 2001; Maamar and Dubnau, 2005). Our observation that *comKS* overexpression restores competence of the *ytrA* mutant only partially suggests that ComK levels are not the only factor that limits competence of the *ytrA* mutant. If the *ytrA* deletion would interfere with ComK activation, one would then expect wild type like competence upon overexpression of ComK which was not the case. Why does ComK then restore the competence at all? The DNA uptake apparatus must be adapted to cell wall thickness in order to ensure that the extracellular DNA can reach the ComG/ComE DNA transport complex. Due to the increased cell wall thickness upon overexpression of the YtrBCDEF ABC transporter, the DNA probably has problems to get in contact with the ComG pili. Overexpression of ComK will then result in the increased production of DNA-binding ComG on the cell surface of all cells of the population (comparing to about 10% in the wild-type strain transformed with the classical two-step protocol). This would simply increase the probability that foreign DNA reaches the DNA uptake machinery in some cells, which then leads to the appearance of only a few transformants as observed in our study. On the other hand, the results obtained by fluorescence microscopy revealed a decreased transcription from the ComK dependent *comG* promoter in the *ytrA* mutant. However, this expression is expected to be wild type-like if the action of YtrBCEDF ABC transporter would not interfere with ComK activity and only block DNA uptake as a result of the remodeled cell wall as suggested above. Again, the disorganized cell wall might be responsible, since ComK expression is induced by the detection of extracellular quorum-sensing signals (both ComXPA and Rap-Phr systems) and this induction depends on the accessibility of the sensor domains for the pheromones which might be impaired in the strain with altered cell wall composition.

In addition to the loss of genetic competence, it was previously shown that the *ytrA* deletion leads to decreased sporulation efficiency (Koo et al, 2017) and we have shown that it also affects biofilm formation. Considering the changed cell wall properties, this is in agreement with previous studies which showed hampered biofilm formation upon disruption of cell wall biosynthesis (Bucher et al., 2015; Zhu et al., 2018). Taken together, we conclude that the overexpression of the YtrBCDEF ABC transporter upon deletion of *ytrA* plays a pleiotropic role in the control of alternative lifestyles of *B. subtilis.*

Our results demonstrate that the YtrBCDEF ABC transporter is involved in the control of cell wall homeostasis, but it is not yet clear how this is achieved. An easy explanation would be that the system exports molecules necessary for cell wall synthesis, however, based on the presence of the solute binding protein YtrF and on the critical role of this protein in preventing genetic competence, it can be assumed that the ABC transporter rather acts as an importer. However, YtrBCDEF may not act as a transporter at all and simply modulate the activity of other enzymes that participate in cell wall metabolism. Strikingly, YtrF is a member of the same protein family as FtsX, which is known to activate the cell wall hydrolase CwlO (Meisner et al., 2013). Future work will need to address the precise mechanism by which the YtrBCDEF ABC transporter interferes with cell wall synthesis.

## ACKNOWLEDGEMENTS

We wish to thank Julia Busse, Melin Güzel and Leon Daniau for the help with some experiments. We are grateful to Josef Altenbuchner, Jan Gundlach, Leendert Hamoen, Daniel Kearns, Daniel Reuss, Sarah Wilcken, and the Bacillus Genetic Sock Center for providing *B. subtilis* strains. We thank Dr. Michael Hoppert for providing access to the Transmission Electron Microscope. This research received funding from the Deutsche Forschungsgemeinschaft via SFB860.

## Notes

### Competing Interest Statement

The authors have declared no competing interest.

## References

Aguilar, C., Vlamakis, H., Guzman, A., Losick, R., and Kolter, R. (2010) KinD is a checkpoint protein linking spore formation to extracellular-matrix production in *Bacillus subtilis* biofilms. mBio 1, e00035–10.

Beckering, C. L., Steil, L., Weber, M. H. W., Völker, U., and Marahiel, M. A. (2002) Genome-wide transcriptional analysis of the cold shock response in *Bacillus subtilis*. J. Bacteriol. 184, 6395–6402.

Berka, R. M., Hahn, J., Albano, M., Draskovic, I., Persuh, M., Cui, X., et al. (2002) Microarray analysis of the *Bacillus subtilis* K-state: Genome-wide expression changes dependent on ComK. Mol. Microbiol. 43, 1331–1345.

Beveridge, T. J., and Murray, R. G. E. (1979) How thick is the *Bacillus subtilis* cell wall? Curr. Microbiol. 2, 1–4.

Boonstra, M., Schaffer, M., Sousa, J., Morawska, L., Holsappel, S.,Hildebrandt, P., et al. (2020) Analyses of competent and non-competent subpopulations of *Bacillus subtilis* reveal *yhfW, yhxC* and ncRNAs as novel players in competence. Environ. Microbiol. 22, 2312–2328.

Branda, S. S., González-Pastor, J. E., Ben-Yehuda, S., Losick, R., and Kolter, R. (2001) Fruiting body formation by *Bacillus subtilis*. Proc. Natl. Acad. Sci. U. S. A. 98, 11621–11626.

Bucher, T., Oppenheimer-Shaanan, Y., Savidor, A., Bloom-Ackermann, Z., and Kolodkin-Gal, I. (2015) Disturbance of the bacterial cell wall specifically interferes with biofilm formation. Environ. Microbiol. Rep. 7, 990–1004.

Cascante-Estepa, N., Gunka, K., and Stülke, J. (2016) Localization of components of the RNA-degrading machine in *Bacillus subtilis*. Front. Microbiol. 7, 1492.

Cao, M., Wang, T., Ye, R., and Helmann, J. D. (2002) Antibiotics that inhibit cell wall biosynthesis induce expression of the *Bacillus subtilis* σ^W^ and σ^M^ regulons. Mol. Microbiol. 45, 1267–1276.

Diethmaier, C., Pietack, N., Gunka, K., Wrede, C., Lehnik-Habrink, M., Herzberg, C., et al. (2011) A novel factor controlling bistability in *Bacillus subtilis:* The YmdB protein affects flagellin expression and biofilm formation. J. Bacteriol. 193, 5997–6007.

Draskovic, I., and Dubnau, D. (2005) Biogenesis of a putative channel protein, ComEC, required for DNA uptake: Membrane topology, oligomerization and formation of disulphide bonds. Mol. Microbiol. 55, 881–896.

Figaro, S., Durand, S., Gilet, L., Cayet, N., Sachse, M., and Condon, C. (2013) *Bacillus subtilis* mutants with knockouts of the genes encoding ribonucleases RNase Y and RNase J1 are viable, with major defects in cell morphology, sporulation, and competence. J. Bacteriol. 195, 2340–2348.

Flórez, L. A., Gunka, K., Polanía, R., Tholen, S., and Stülke, J. (2011) SPABBATS: A pathway-discovery method based on Boolean satisfiability that facilitates the characterization of suppressor mutants. BMC Syst. Biol. 5, 5.

Gamba, P., Jonker, M. J., and Hamoen, L. W. (2015) A novel feedback loop that controls bimodal expression of genetic competence. PLoS Genet. 11, e1005047.

Guérout-Fleury, A. M., Shazand, K., Frandsen, N., and Stragier, P. (1995) Antibiotic-resistance cassettes for *Bacillus subtilis*. Gene 167, 335–336.

Hahn, J., Roggiani, M., and Dubnau, D. (1995) The major role of Spo0A in genetic competence is to downregulate *abrB*, an essential competence gene. J. Bacteriol. 195, 177, 3601–3605.

Haijema, B. J., Hahn, J., Haynes, J., and Dubnau, D. (2001) A ComGA-dependent checkpoint limits growth during the escape from competence. Mol. Microbiol. 40, 52–64.

Hamoen, L. W., Kausche, D., Marahiel, M. A., Sinderen, D., Venema, G., and Serror, P. (2003) The *Bacillus subtilis* transition state regulator AbrB binds to the −35 promoter region of *comK*. FEMS Microbiol. Lett. 218, 299–304.

Hamoen, L. W., Smits, W. K., de Jong, A., Holsappel, S., and Kuipers, O. P. (2002) Improving the predictive value of the competence transcription factor (ComK) binding site in *Bacillus subtilis* using a genomic approach. Nucleic Acids Res. 30, 5517–5528.

Hamoen, L. W., Van Werkhoven, A. F., Venema, G., and Dubnau, D. (2000) The pleiotropic response regulator DegU functions as a priming protein in competence development in *Bacillus subtilis*. Proc. Natl. Acad. Sci. U. S. A. 97, 9246–9251.

Hoa, T. T., Tortosa, P., Albano, M., and Dubnau, D. (2002) Rok (YkuW) regulates genetic competence in *Bacillus subtilis* by directly repressing *comK*. Mol. Microbiol. 43, 15–26.

Hutter, B., Fischer, C., Jacobi, A., Schaab, C., and Loferer, H. (2004) Panel of *Bacillus subtilis* reporter strains indicative of various modes of action. Antimicrob. Agents Chemother. 48, 2588–2594.

Kampf, J., Gerwig, J., Kruse, K., Cleverley, R., Dormeyer, M., Grünberger, A., et al. (2018) Selective pressure for biofilm formation in *Bacillus subtilis:* Differential effect of mutations in the master regulator SinR on bistability. mBio 9, e01464–18.

Konkol, M. A., Blair, K. M., and Kearns, D. B. (2013) Plasmid-encoded ComI inhibits competence in the ancestral 3610 strain of *Bacillus subtilis*. J. Bacteriol. 195, 4085–4093.

Koo, B. M., Kritikos, G., Farelli, J. D., Todor, H., Tong, K., Kimsey, H., et al. (2017) Construction and analysis of two genome-scale deletion libraries for *Bacillus subtilis*. Cell Syst. 4, 291–305.

Kunst, F., and Rapoport, G. (1995) Salt stress is an environmental signal affecting degradative enzyme synthesis in *Bacillus subtilis*. J. Bacteriol. 177, 2403–2407.

López, D., Vlamakis, H., and Kolter, R. (2009) Generation of multiple cell types in *Bacillus subtilis*. FEMS Microbiol. Rev. 33, 152–163.

López, D., and Kolter, R. (2010) Extracellular signals that define distinct and coexisting cell fates in *Bacillus subtilis*. FEMS Microbiol. Rev. 34, 134–149.

Luttinger, A., Hahn, J., and Dubnau, D. (1996) Polynucleotide phosphorylase is necessary for competence development in *Bacillus subtilis*. Mol. Microbiol. 19, 343–356.

Maamar, H., and Dubnau, D. (2005) Bistability in the *Bacillus subtilis* K-state (competence) system requires a positive feedback loop. Mol. Microbiol. 56, 615–624.

Maier, B. (2020) Competence and transformation in *Bacillus subtilis*. Curr. Issues Mol. Biol. 37, 57–76.

Mascher, T., Margulis, N. G., Wang, T., Ye, R. W., and Helmann, J. D. (2003) Cell wall stress responses in *Bacillus subtilis:* The regulatory network of the bacitracin stimulon. Mol. Microbiol. 50, 1591–1604.

Mechold, U., Fang, G., Ngo, S., Ogryzko, V., and Danchin, A. (2007) YtqI from *Bacillus subtilis* has both oligoribonuclease and pAp-phosphatase activity. Nucleic Acids Res. 35, 4552–4561.

Meisner, J., Montero Llopis, P., Sham, L.-T., Garner, E., Bernhardt, T. G., and Rudner, D. Z. (2013) FtsEX is required for CwlO peptidoglycan hydrolase activity during cell wall elongation in *Bacillus subtilis*. Mol. Microbiol. 89, 1069–1083.

Mirouze, N., Desai, Y., Raj, A., and Dubnau, D. (2012) Spo0A~P imposes a temporal gate for the bimodal expression of competence in *Bacillus subtilis*. PLoS Genet. 8, e1002586.

Mirouze, N., Ferret, C., Cornilleau, C., and Carballido-López, R. (2018) Antibiotic sensitivity reveals that wall teichoic acids mediate DNA binding during competence in *Bacillus subtilis*. Nat. Commun. 9, 5072.

Nakano, M. M., Xia, L., and Zuber, P. (1991). Transcription initiation region of the *srfA* operon, which is controlled by the *comP-comA* signal transduction system in *Bacillus subtilis*. J. Bacteriol. 173, 5487–5493.

Nicolas, P., Mäder, U., Dervyn, E., Rochat, T., Leduc, A., Pigeonneau, N., et al. (2012) The condition-dependent whole-transcriptome reveals high-level regulatory architecture in bacteria. Science 335, 1103–1106.

Ogura, M., Yamaguchi, H., Kobayashi, K., Ogasawara, N., Fujita, Y., and Tanaka, T. (2002) Whole-genome analysis of genes regulated by the *Bacillus subtilis* competence transcription factor ComK. J. Bacteriol. 184, 2344–2351.

Quentin, Y., Fichant, G., and Denizot, F. (1999) Inventory, assembly and analysis of *Bacillus subtilis* ABC transport systems. J. Mol. Biol. 287, 467–484.

Rahmer, R., Heravi, K. M., and Altenbuchner, J. (2015) Construction of a super-competent *Bacillus subtilis* 168 using the *PmtlA-comKS* inducible cassette. Front. Microbiol. 6, 1431.

Reuß, D. R., Altenbuchner, J., Mäder, U., Rath, H., Ischebeck, T., Sappa, P. K., et al. (2017) Large-scale reduction of the *Bacillus subtilis* genome: Consequences for the transcriptional network, resource allocation, and metabolism. Genome Res. 27, 289–299.

Rincón Tomás, B., González, F. J., Somoza, L., Sauter, K., Madureira, P., Medialdea, T., et al. (2020) Siboglinidae tubes as an additional niche for microbial communities in the Gulf of Cádiz – A microscopical appraisal. Microorganisms 8, 367.

Rueden, C. T., Schindelin, J., Hiner, M. C., DeZonia, B. E., Walter, A. E., Arena, E. T., and Eliceiri, K. W. (2017) ImageJ2: ImageJ for the next generation of scientific image data, BMC Bioinformatics 18, 529.

Salzberg, L. I., Luo, Y., Hachmann, A.-B., Mascher, T., and Helmann, J. D. (2011) The *Bacillus subtilis* GntR family repressor YtrA responds to cell wall antibiotics. J. Bacteriol. 193, 5793–5801.

Sambrook, J., Fritsch, E. F., and Maniatis, T. (1989) Molecular cloning: a laboratory manual, 2nd ed. Cold Spring Harbor Laboratory, Cold Spring Harbor, N.Y. 1989

Serror, P., and Sonenshein, A. L. (1996) CodY is required for nutritional repression of *Bacillus subtilis* genetic competence. J. Bacteriol. 178, 5910–5915.

Shimane, K., and Ogura, M. (2004) Mutational analysis of the helix-turn-helix region of *Bacillus subtilis* response regulator DegU, and identification of *cis*-acting sequences for DegU in the *aprE* and *comK* promoters. J. Biochem. 136, 387–397.

Smits, W. K., Eschevins, C. C., Susanna, K. A., Bron, S., Kuipers, O. P., and Hamoen, L. W. (2005) Stripping *Bacillus:* ComK auto-stimulation is responsible for the bistable response in competence development. Mol. Microbiol. 56, 604–614.

Turgay, K., Hahn, J., Burghoorn, J., and Dubnau, D. (1998) Competence in *Bacillus subtilis* is controlled by regulated proteolysis of a transcription factor. EMBO J. 17, 6730–6738.

van Sinderen, D., Luttinger, A., Kong, L., Dubnau, D., Venema, G., and Hamoen, L. (1995) *comK* encodes the competence transcription factor, the key regulatory protein for competence development in *Bacillus subtilis*. Mol. Microbiol. 15, 455–462.

Wach, A. (1996) PCR-synthesis of marker cassettes with long flanking homology regions for gene disruptions in *S. cerevisiae*. Yeast 12, 259–265.

Wenzel, M., Kohl, B., Münch, D., Raatschen, N., Albada, H. B., Hamoen, L., et al. (2012) Proteomic response of *Bacillus subtilis* to lantibiotics reflects differences in interaction with the cytoplasmic membrane. Antimicrob. Agents Chemother. 56, 5749–5757.

Yoshida, K.-I., Fujita, Y., and Dusko Ehrlich, A. S. (2000) An operon for a putative ATP-binding cassette transport system involved in acetoin utilization of *Bacillus subtilis*. J. Bacteriol. 182, 5454–5461.

Youngman, P. (1990) Use of transposons and integrational vectors for mutagenesis and construction of gene fusions in *Bacillus subtilis*. Mol Biol Methods for Bacillus, 221–266.

Zhu, X., Liu, D., Singh, A. K., Drolia, R., Bai, X., Tenguria, S., and Bhunia, A. K. (2018) Tunicamycin mediated inhibition of wall teichoic acid affects *Staphylococcus aureus* and *Listeria monocytogenes* cell morphology, biofilm formation and virulence. Front. Microbiol. 9, 1352.

