## Supplementary material for "The YtrBCDEF ABC transporter is involved in the control of social activities in *Bacillus subtilis*": S1

TABLE S1. Oligonucleotides used in this study.

| **Primer** | **Sequence ^a^** | **Purpose** |
| --- | --- | --- |
| VK17 | GACGAAGACGGAAATGAGCTAGATGC | fwd; Amplification of upstream fragment (deletion of *greA*) |
| VK18 | *CCTATCACCTCAAATGGTTCGCTG*GTTCAAGTTTTTGTTTTCCTTCTGCAGTCATAGG | rev; Amplification of upstream fragment (deletion of *greA*) |
| VK19 | *CGAGCGCCTACGAGGAATTTGTATCG*GATGAAGAAGTCACAGTACAAACACCGG | fwd; Amplification of downstream fragment (deletion of *greA*) |
| VK20 | TGCAGCTGCGGCAATGACTGTTTTAAAAAC | rev; Amplification of downstream fragment (deletion of *greA*) |
| VK21 | GGCTTAGTGCTGAATTATGATGAAGATACAC | fwd; Sequencing *greA* deletion |
| VK22 | GTGCCTTTGTCGTCCCCCGG | rev; Sequencing *greA* deletion |
| JN420 | *CCTATCACCTCAAATGGTTCGCTG*CGCACATGTCTATGTAAGATAATCGT | rev; Amplification of upstream fragment (deletion of *nrnA*) |
| JN421 | GGGATCGAAGTGCTTCCCG | fwd; Amplification of upstream fragment (deletion of *nrnA*) |
| JN422 | *CCGAGCGCCTACGAGGAATTTGTATCG*GCTGGGATGAAGCTGATCGTA | fwd; Amplification of downstream fragment (deletion of *nrnA*) |
| JN423 | GCGGCATACTCGAAGGCA | rev; Amplification of downstream fragment (deletion of *nrnA*) |
| JN424 | GACCAAAAATCCCGTCACGG | fwd; Sequencing *nrnA* deletion |
| JN425 | GCTTGCCAACCGGTTAAAAATATG | rev; Sequencing *nrnA* deletion |
| MB31 | CTGCGTATATCTGCTTCGAAATCCTTC | fwd; Amplification of upstream fragment (integration of P*_mtlA_-comKS*) |
| MB32 | *TAAAAATAAAAAGCTAGCGGGGATCCCAAG*TCAAAACCGAGTCTCATTTCCTATTTATCC | rev; Amplification of upstream fragment (integration of P*_mtlA_-comKS*) |
| MB33 | CTTGGGATCCCCGCTAGCTTTTTATTTTTA | fwd; Amplification of P*_mtlA_-comKS* for its insertion into *yvcA-hisI* locus |
| MB34 | *CCTATCACCTCAAATGGTTCGCTG*CGGAGGATTTCGTGCCGGTTGATTA | rev; Amplification of P*_mtlA_-comKS* for its insertion into *yvcA-hisI* locus |
| MB35 | *CCGAGCGCCTACGAGGAATTTGTATCG* GCCGGCTAGCACCCAATATAAATCTAAAT | fwd; Amplification of downstream fragment (integration of P*_mtlA_-comKS*) |
| MB36 | GTGCTGACACTTGCGTATATGAACAAG | rev; Amplification of downstream fragment (integration of P*_mtlA_-comKS*) |
| MB37 | GTAAACTCCTTTGTAGCCTCATACTGAC | fwd; Sequencing P*_mtlA_-comKS* insertion |
| MB38 | GAATGTGAGATGAAACAGGCAGATGAAC | rev; Sequencing P*_mtlA_-comKS* insertion |
| MB43 | CTTGATAGATACTTTCCATCCTCCGG | fwd; Sequencing P*_mtlA_-comKS* insertion |
| MB44 | CCCTACACTTTCTTCGACAAGACCC | fwd; Sequencing P*_mtlA_-comKS* insertion |
| MB60 | GCTGATGAAACGGCAGTGCT | fwd; Amplification of upstream fragment (deletion of *ftsH*) |
| MB61 | *CCTATCACCTCAAATGGTTCGCTG*TCCTTACCTCCTCCCACAG | rev; Amplification of upstream fragment (deletion of *ftsH*) |
| MB62 | *CCGAGCGCCTACGAGGAATTTGTATCG*AAGACGATACGAAAGAGTAATTCGC | fwd; Amplification of downstream fragment (deletion of *ftsH*) |
| MB63 | CTCCTATACACTTCCTACGCGG | rev; Amplification of downstream fragment (deletion of *ftsH*) |
| MB64 | GGGCTGAAGGTGGTCAAATC | fwd; Sequencing *ftsH* deletion |
| MB65 | CATATCAGTCGTTCTCGCTGCA | rev; Sequencing *ftsH* deletion |
| MB66 | CATCGGTCCGGTTTCCAGCA | fwd; Amplification of upstream fragment (deletion of *ytrA*) |
| MB67 | *CCTATCACCTCAAATGGTTCGCTG*GGGTGTTGAGCTTCTTGGATC | rev; Amplification of upstream fragment (deletion of *ytrA*) |
| MB68 | *CCGAGCGCCTACGAGGAATTTGTATCG*GCTGATGTGAAGGGAGGCAA | fwd; Amplification of downstream fragment (deletion of *ytrA*) |
| MB69 | GGCGATCAAGACACCCTTGA | rev; Amplification of downstream fragment (deletion of *ytrA*) |
| MB70 | GATGTACTTGCCGTCCTTCCA | fwd; Sequencing *ytrA* deletion |
| MB71 | ACCCGGCACCCAGTTGATAT | rev; Sequencing *ytrA* deletion |
| MB72 | AGGGGACAGAGTATCTCAGGCA | fwd; Amplification of upstream fragment (deletion of *comEC*) |
| MB73 | *CCTATCACCTCAAATGGTTCGCTG*CGCATTCATCACACGTAGCTC | rev; Amplification of upstream fragment (deletion of *comEC*) |
| MB74 | *CCGAGCGCCTACGAGGAATTTGTATCG*AAAGACTGCCGAGAAATCAGCA | fwd; Amplification of downstream fragment (deletion of *comEC*) |
| MB75 | TCTCCAATAAACGTGCAGAGCTT | rev; Amplification of downstream fragment (deletion of *comEC*) |
| MB76 | AACAACGACGAGTCAAACGAAACAA | fwd; Sequencing *comEC* deletion |
| MB77 | CTCTGTTCGTTTTCGGTTGACG | rev; Sequencing *comEC* deletion |
| MB78 | AACCGTTTATCCGAGGTCAGCC | fwd; Amplification of upstream fragment (deletion of *degU*) |
| MB79 | *CCTATCACCTCAAATGGTTCGCTG*GTTTACTTTAGTCACAAGCCACGC | rev; Amplification of upstream fragment (deletion of *degU*) |
| MB80 | *CCGAGCGCCTACGAGGAATTTGTATCG*GGCTGGGTAGAAATGAGATAGTA | fwd; Amplification of downstream fragment (deletion of *degU*) |
| MB81 | AGCACGCCTCCTTTCGAAACAG | rev; Amplification of downstream fragment (deletion of *degU*) |
| MB82 | GCAGGTGTATGAAGTGATTGAGC | fwd; Sequencing *degU* deletion |
| MB83 | TCGAAGCGTCTGCTGCAATTC | rev; Sequencing *degU* deletion |
| MB70 | GATGTACTTGCCGTCCTTCCA | fwd; Amplification of upstream fragment (deletion of *ytr* operon) |
| MB118 | *CCTATCACCTCAAATGGTTCGCTG*CACTTAATACAATAAATACTTTGACTCACA | rev; Amplification of upstream fragment (deletion of *ytr* operon) |
| MB119 | *CCGAGCGCCTACGAGGAATTTGTATCG*TATAATGCGAACGAGCCGGC | fwd; Amplification of downstream fragment (deletion of *ytr* operon) |
| MB120 | GCACAAATACACCATATAAAGTACATTCC | rev; Amplification of downstream fragment (deletion of *ytr* operon) |
| MB121 | CGATCGAAATGCCGACCAC | fwd; Sequencing *ytr* operon deletion |
| MB122 | GTTCATTTATGGCTGTCACATCGAG | rev; Sequencing *ytr* operon deletion |
| MB70 | GATGTACTTGCCGTCCTTCCA | fwd; Amplification of upstream fragment (deletion of *ytrG-E* region) |
| MB118 | *CCTATCACCTCAAATGGTTCGCTG*CACTTAATACAATAAATACTTTGACTCACA | rev; Amplification of upstream fragment (deletion of *ytrG-E* region) |
| MB194 | *CCGAGCGCCTACGAGGAATTTGTATCG*TTGAGGTTTAAGGATCAGGTTCATTTTAT | fwd; Amplification of downstream fragment (deletion of *ytrG-E* region) |
| MB195 | GATACATCCGACAAAGATCAGTCC | rev; Amplification of downstream fragment (deletion of *ytrG-E* region) |
| MB121 | CGATCGAAATGCCGACCAC | fwd; Sequencing *ytrG-E* region deletion |
| MB187 | TTATAATTCTCTTCTCAACGCTGTCAG | rev; Sequencing *ytrG-E* region deletion |
| MB68 | *CCGAGCGCCTACGAGGAATTTGTATCG*GCTGATGTGAAGGGAGGCAA | fwd; Confirmation of *ytrC* deletion |
| MB180 | GACACAGCCTTGATAGATGAGATAC | rev; Confirmation of *ytrC* deletion |
| CB449 | *CCTATCACCTCAAATGGTTCGCTG*TCCTTAAACCTCAACGGTAATTCCT | fwd; Confirmation of *ytrD* deletion |
| MB179 | CTGGATTCTTTGTGAGCTACTTCTC | rev; Confirmation of *ytrD* deletion |
| CB448 | TCACCATATTATTTAGTCATTCCGGC | fwd; Confirmation of *ytrE* deletion |
| CB449 | *CCTATCACCTCAAATGGTTCGCTG*TCCTTAAACCTCAACGGTAATTCCT | rev; Confirmation of *ytrE* deletion |
| MB121 | CGATCGAAATGCCGACCAC | Fwd; Amplification of *ytrAB:ermC* from GP3193 |
| MB180 | GACACAGCCTTGATAGATGAGATAC | rev; Amplification of *ytrAB:ermC* from GP3193 |
| MB186 | TTTTCTAGATATGAGGTTTAAGGATCAGGTTCATTTTAT | fwd; Amplification of *ytrF* for cloning into pGP888, XbaI |
| MB187 | ATTGGTACCTTATAATTCTCTTCTCAACGCTGTCAG | rev; Amplification of *ytrF* for cloning into pGP888, KpnI |
| MB198 | GCGGCAGCTGTCAAAAGC | fwd; Amplification of upstream fragment (deletion of *ytrCD*) |
| MB199 | *CCTATCACCTCAAATGGTTCGCTG*CTACCATCTCCGCTTCCCTC | rev; Amplification of upstream fragment (deletion of *ytrCD*) |
| MB200 | *CCGAGCGCCTACGAGGAATTTGTATCG*GTAAGGGAGAGAGAACATATGATTG | fwd; Amplification of downstream fragment (deletion of *ytrCD*) |
| MB201 | CTCCTTCCTTGCCCATTACG | rev; Amplification of downstream fragment (deletion of *ytrCD*) |
| MB202 | CACTATGCAGGGGTTGAGCT | fwd; Sequencing *ytrCD* deletion |
| MB203 | GTTTGGTTCATACACTTGCGTTC | rev; Sequencing *ytrCD* deletion |
| CB448 | TCACCATATTATTTAGTCATTCCGGC | fwd; Amplification of upstream fragment (deletion of *ytrF*) |
| CB449 | *CCTATCACCTCAAATGGTTCGCTG*TCCTTAAACCTCAACGGTAATTCCT | rev; Amplification of upstream fragment (deletion of *ytrF*) |
| CB450 | *CGAGCGCCTACGAGGAATTTGTATCG*TTATAATGCGAACGAGCCGGCT | fwd; Amplification of downstream fragment (deletion of *ytrF*) |
| CB451 | TCCCATGTTTTCAAGCTTTTATAAAACG | rev; Amplification of downstream fragment (deletion of *ytrF*) |
| CB452 | ACCTCGAGATCCTTTTTGGCG | fwd; Sequencing *ytrF* deletion |
| CB453 | TGCTAAGCGATGCCGTGCT | rev; Sequencing *ytrF* deletion |
| cat-fwd (kan) | *CAGCGAACCATTTGAGGTGATAGG*CGGCAATAGTTACCCTTATTATCAAG | fwd; Amplification of chloramphenicol resistance cassette |
| cat-rev (kan) | *CGATACAAATTCCTCGTAGGCGCTCGG*CCAGCGTGGACCGGCGAGGCTAGTTACCC | rev; Amplification of chloramphenicol resistance cassette |
| cat-rev w/o T. (kan) | *CGATACAAATTCCTCGTAGGCGCTCGG*TTATAAAAGCCAGTCATTAGGCCTATC | rev; Amplification of chloramphenicol resistance cassette without Term. |
| kan-fwd | CAGCGAACCATTTGAGGTGATAGG | fwd; Amplification of kanamycin resistance cassette |
| kan-rev | CGATACAAATTCCTCGTAGGCGCTCGG | rev; Amplification of kanamycin resistance cassette |
| mls-fwd (kan) | *CAGCGAACCATTTGAGGTGATAGG*GATCCTTTAACTCTGGCAACCCTC | fwd; Amplification of erythromycin resistance cassette |
| mls-rev (kan) | *CGATACAAATTCCTCGTAGGCGCTCGG*GCCGACTGCGCAAAAGACATAATCG | rev; Amplification of erythromycin resistance cassette |
| mls-rev w/o T. (kan) | *CGATACAAATTCCTCGTAGGCGCTCGG*TTACTTATTAAATAATTTATAGCTATTG | rev; Amplification of erythromycin resistance cassette without Term. |
| spc-fwd (kan) | *CAGCGAACCATTTGAGGTGATAGG*GACTGGCTCGCTAATAACGTAACGTGACTGGCAAGAG | fwd; Amplification of spectinomycin resistance cassette |
| spc-rev w/o T. (kan) | *CGATACAAATTCCTCGTAGGCGCTCGG*GTAGTATTTTTTGAGAAGATCAC | rev; Amplification of spectinomycin resistance cassette without Term. |

^a^ Homologous bases for joining PCR are shown in italics, restriction sites are underlined.
